# Neural Computational Model Predicts Attentional Dynamics during Immersive Search in Virtual Reality

**DOI:** 10.1101/2025.10.07.680957

**Authors:** Joyce Tam, Chloe Callahan-Flintoft

## Abstract

Much of the scientific understanding of visual attention has come from desktop paradigms where body, head, and eye movements are restricted. This stands in contrast to the ability to search and navigate with few constraints in the real world. To bridge this gap, a computational model parametrized on a wide range of well-controlled desktop search tasks was evaluated on its ability to predict search behaviors in a less constrained, immersive environment in virtual reality. A set of validation metrics showed that the model’s attention map could predict empirical behaviors such as eye gaze and manual responses in VR despite the field-of-view varying from one moment to another from the participant’s movements. The present work quantifies the real-world applicability of laboratory-based theory and highlights ways to address outstanding limitations in explaining naturalistic visual search behaviors.

---

At any given moment, we are presented with a vast amount of visual information, deeming it necessary for our brain to perpetually parse the visual world by prioritizing selective subsets of information – a collection of mechanisms referred to as attention^1^. Despite the theorized importance of visual attention in supporting everyday tasks such as search and navigation, much of its scientific understanding is based on strictly controlled laboratory studies where simplistic visual arrays are viewed from a restricted vantage point^2–4^. It is therefore unclear how well theories built on this type of laboratory experimentation can explain visual processing in the real world which concerns three-dimensional and dynamic information that is navigated with few constraints^5^.

The first challenge in bridging the gap between laboratory studies used to isolate mechanisms of visual attention and real-world tasks in which those mechanisms are used is being able to perform controlled experimentation in three-dimensional, immersive environments where participants can move as they could in the real world. Virtual reality (VR) offers a solution to this challenge where much of the immersion and interactions available in the real world is simulated while control over spatiotemporal parameters of visual elements is maintained^6^. Additionally, high sampling rate eye trackers and accelerometers can be incorporated to obtain precise measures of gaze, head, and body movement^7,8^. Studies taking advantage of VR, while still relatively rare, have provided important insights into the real-world applicability (or lack thereof) of attention phenomena observed in laboratory paradigms^9–11^.

The second major challenge in bridging the gap between the lab and the real world is being able to test the sheer number of mechanisms and phenomena discovered using desktop paradigms in naturalistic vision. The literature has documented a wide variety of behavioral phenomenon with linkage to different enhancement^12^ and inhibitory^13^ mechanisms, all thought to be encompassed under the term “visual attention”. Validating each of these findings individually in an ecologically valid context would be extremely costly and time-consuming. The solution proposed here is to use a computational model, built and parameterized on a wealth of empirical desktop paradigm findings, to test whether the collection of mechanisms embodied in the model can predict attentional deployment in a more immersive, less constrained environment.

The computational model used is called Reflexive Attention Gradient through Neural AttRactOr Competition^24^ (RAGNAROC; Figure 1). RAGNAROC is a rate coded neural model which simulates the spatiotemporal deployment of covert, reflexive attention. The two adjustable parameters are “bottom-up” saliency weightings of the presented stimuli which represent visual conspicuity^25^, and the “top-down” relevance weightings which represent the importance of specific visual features^26^. Saliency and relevance weightings together encapsulate two major facets of attentional modulations^3,27^. In the model, attentional priority is computed as visual information propagates through a hierarchy of retinotopically organized artificial neurons which ultimately reaches the Attention Map (AM). Along the propagation, the modelled neural activity is modulated by both saliency and relevance parameters in combination with spatiotemporal mechanics that govern neural competition dynamics. The AM therefore emerges as a set of dynamic, retinotopic, two-dimensional maps that simulate the allocation of reflexive, covert attention across the visual field, and how it evolves in time. Based on AM dynamics, it can then be determined what locations in the visual field are expected to “win” the competition for attention and what locations “lose” and receive inhibition.

**Figure 1.**
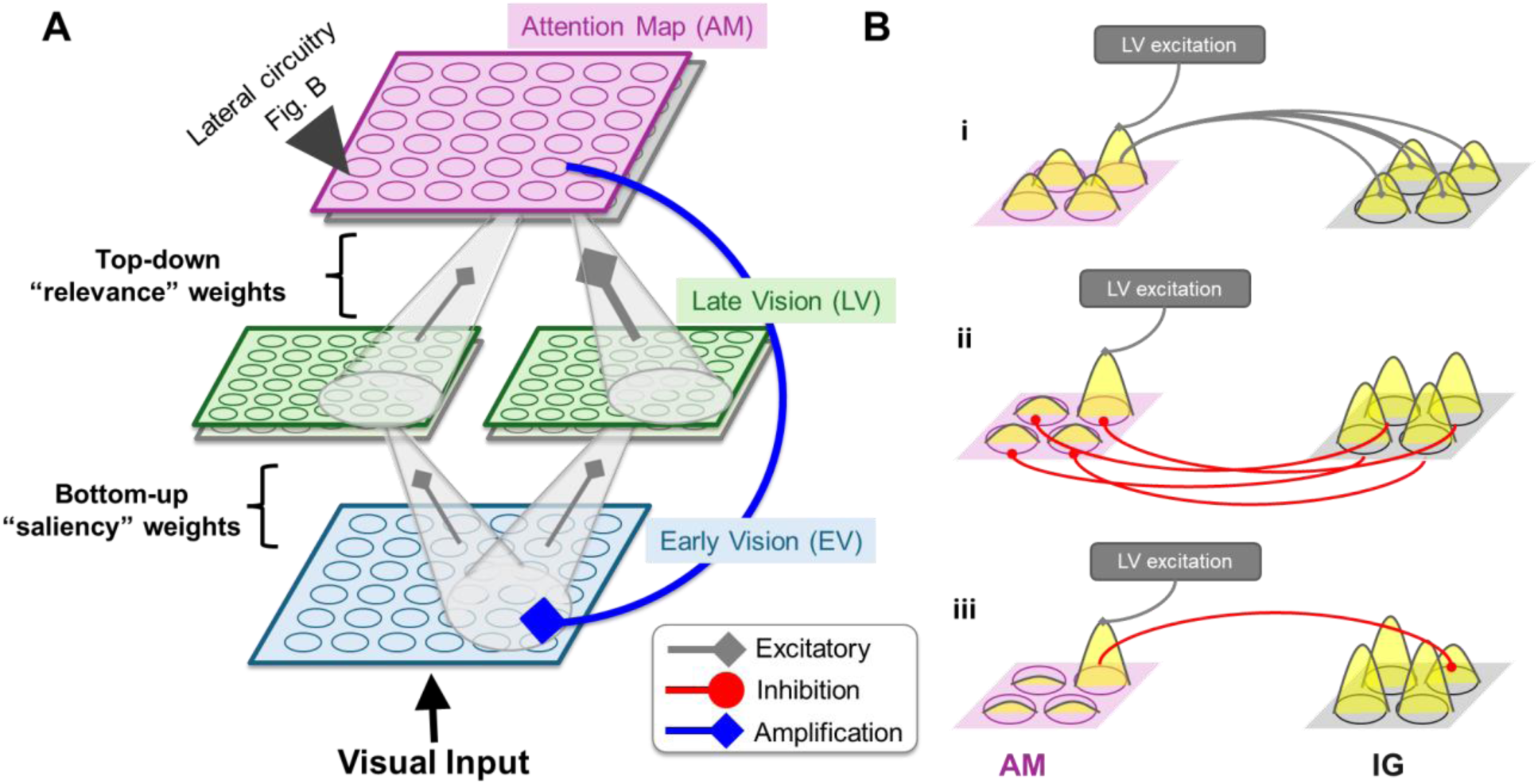
Overview of RAGNAROC’s architecture and lateral circuitry on the Attention Map. (A) Visual input propagates through a hierarchy of neuron maps, during which signals are modulated by bottom-up “saliency” weights and top-down “relevance” weights. Incorporating the influence of both weightings, priority signals on the Attention Map (AM) simulate the attentional landscape across the visual field at a given moment. (B) AM neurons engage in lateral inhibition through Inhibitory Gating (IG) nodes. The inhibitory dynamics allows the model to simulate surround inhibition around an attended location. Step i: An excited AM node excites neighboring IG nodes through a difference-of-Gaussian mask. Step ii: Each IG node inhibits its partnering AM node (one-to-one mapping). Step iii: When an AM node crosses a high threshold, it inhibits its partnering IG node (one-to-one mapping).

RAGNAROC was parametrized upon an array of desktop paradigms to simulate measures of visual search accuracy, reaction time, and electrophysiological data observed in a range of experimental conditions. As such, RAGNAROC encapsulates the state-of-the-art understanding of covert visual attention which ecological validity can thereby be tested as a whole. This modeling work is not dissimilar to other computational models that aim to explain naturalistic visual behaviors, e.g., models to simulate eye gaze patterns during free-viewing^28,29^ or goal-directed search^30,31^ of naturalistic images. However, unlike other models that are trained and tested on data from similar experimental conditions, RAGNAROC was developed to simulate desktop experimentation but now evaluated upon behaviors in an immersive and interactive setting. While this was expected to come at a cost of the model’s predictive accuracy, the success of model simulation in the current work would be uniquely *translational*, indicating the generalizability of laboratory-based theories in real-world contexts.

Additionally, RAGNAROC’s simulated behavioral outcomes in visual search are the product of well-studied and theorized cognitive mechanisms (e.g. attentional enhancement^12,32^, inhibitory surround^33^, reactive inhibition^34^ etc.). The model architecture also references the properties of ventral visual cortical areas^35^ as the processing of visual information and emergence of visual attention was simulated with a hierarchy of neuron maps. Similar to previous models of visual attention that emphasizes biological plausibility^36,37^, the current work not only evaluates the model’s predictive capability but also informs and progresses the understanding of neural attentional mechanisms. This complements alternative modeling approaches that achieve higher predictive accuracy but employ mechanisms that are relatively opaque (e.g., deep neural networks^28,38^) or abstract (e.g., Bayesian inferences^39,40^). As such, the current effort is not only to simulate visual behavior but to test neurocognitive theories in a more ecologically valid environment.

In the current work, participants performed a visual foraging task in VR with a head mounted display. Participants searched for targets defined by a specific color-shape combination among distractors similar to classic conjunction search tasks^41–43^ but with less restrictive eye, head, and body movements (Figure 2).

**Figure 2.**
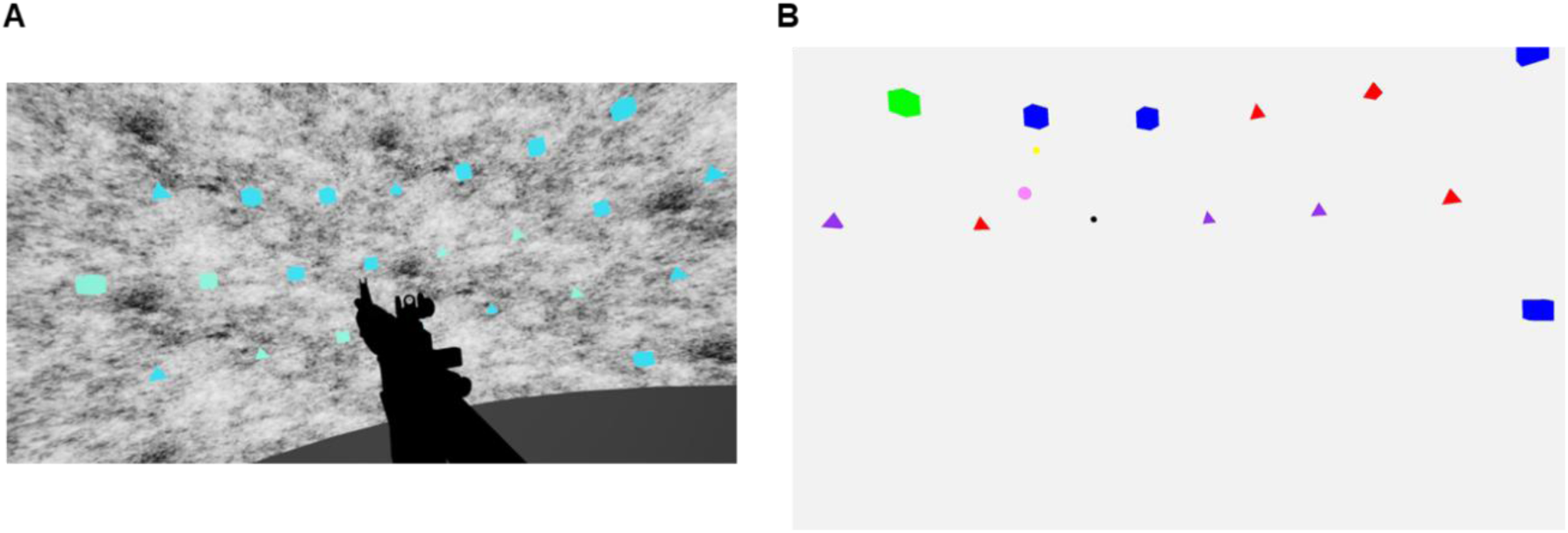
Paradigm and analyzed images. **(A)** An example of the participant’s field-of-view during the experiment. **(B)** An example of a two-dimensional FOV projection that was used to create model input. The two-dimensional projections were in head-centered coordinates which were re-created after the experiment by cross-referencing the moment-to-moment recorded head orientation and stimulus positions in the virtual world. Different stimulus types were coded with arbitrary colors that do not reflect the actual stimulus colors used in the experiment (green = green cubes, blue = blue cubes, red = green pyramids, purple = blue pyramids, pink = probe).

The participant’s field-of-view was sampled at 100Hz to be entered as input to RAGNAROC. The model then output a series of simulated AMs which indicated the predicted attentional priority for each x, y location in the visual field for every timestep. Fixed parameters of the model were unaltered, although two additional parameters were fitted to account for the continuous nature of the input. Specifically, the Model-Data Delay (MDD) characterized the time delay between the input timestep and the predicted timestep and the Gaze Shift Threshold (GST) defined the magnitude of viewpoint change which when crossed, model activation history in retinotopic reference would not be carried over. MDD and GST were fitted on the individual level using a subset of trials before all trials were modelled using the selected parameters.

As RAGNAROC’s simulation of attentional deployment consists of a 2D map which evolves over time, validating such a simulation in its entirety (*x* by *y* by *t* for each participant) is not practical. Instead, a variety of metrics linked to covert and overt attentional deployment were used as proxies to validate model simulation, which included the model’s ability to predict the participants’ moment-to-moment gaze location, the item that was selected for the next fixation or manual response, as well as the reaction time (RT) in reporting the presence of a sudden onset stimulus (see Methods). To preface, evidence to support RAGNAROC’s validity in simulating attentional behaviors in VR was found across all metrics.

While the presented metrics are by no means exhaustive, the current work shows preliminary evidence that RAGNAROC, a computational model grounded in visual attention and search theory and built upon findings from desktop display paradigms, can partially account for overt and covert attentional behaviors of humans in an immersive, unconstrained search task. As such, it could be inferred that the general understanding of visual attention in the field, as constructed through numerous well-controlled experimental studies and now encapsulated within the model, was not confined within walls of the laboratory, and the extent of this generalizability can be effectively quantified through a modeling approach. Overall, our work demonstrates that scientific knowledge in this field has accumulated towards this point where behaviors in day-to-day active vision can be meaningfully predicted and explained, and tangible efforts can be made to further this progress.

## Results

### Adaptive Individual Parameters

The combination of MDD and GST that yielded the lowest mean gaze prediction error in the subsample of trials was chosen for each participant. The mean chosen MDD was 62.30 ms (mode = 0 ms, SD = 220.28 ms), suggesting that gaze prediction error was minimized when a short delay was inserted between the input timestep (based on which the model computed its predictions) and the predicted timestep (upon which the model’s predictions were validated). The mean chosen GST was 0.71 cm (mode = 0 cm, SD = 1.63 cm), meaning that gaze error was smallest when even a small gaze shift led to the complete eradication of model activation history. Figure 3 shows how the mean gaze prediction error varied by MDD and GST.

**Figure 3.**
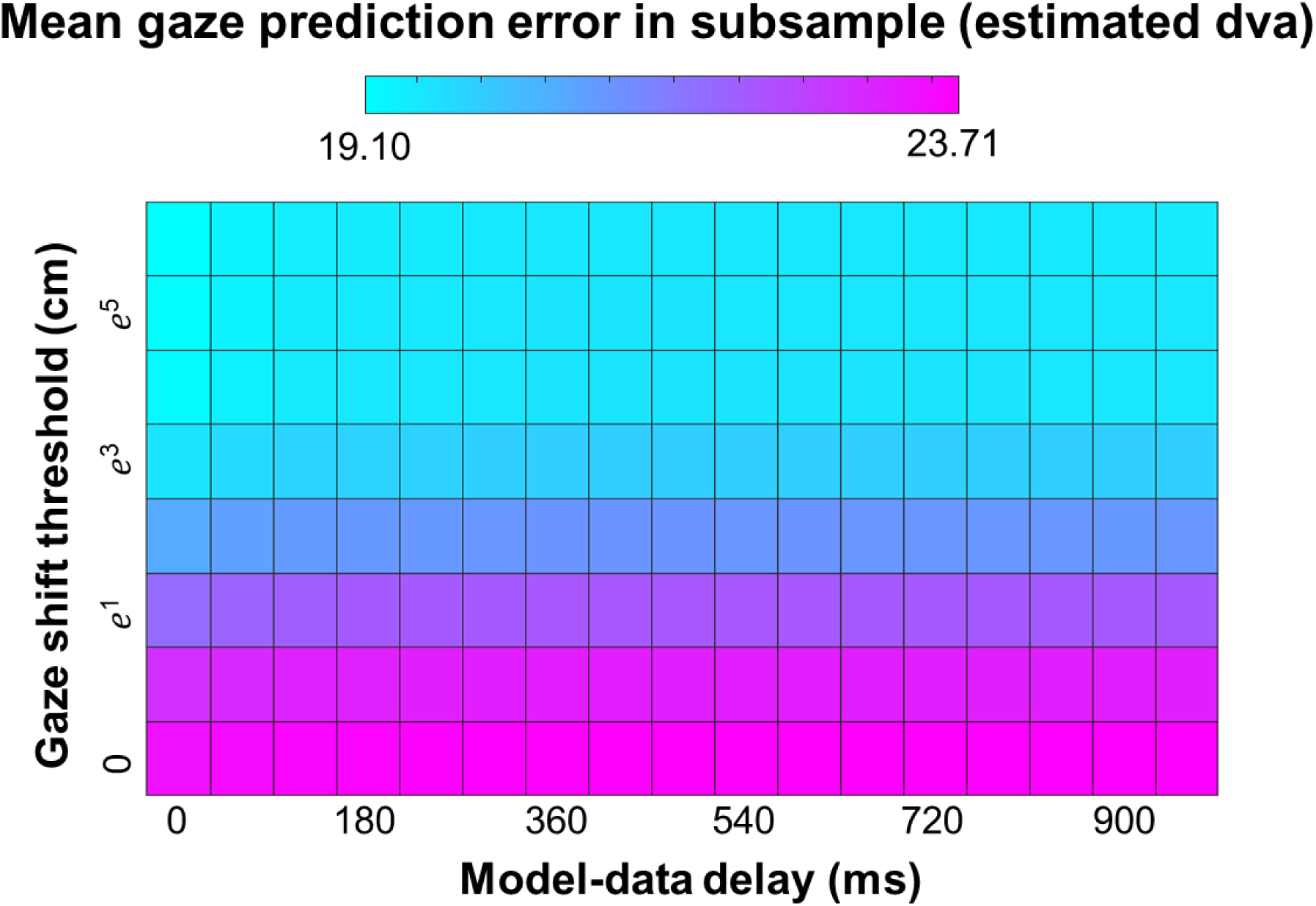
Gaze prediction error in trial subsamples, averaged across participants. Error values varied across model-data delay (MDD; x-axis) and gaze shift threshold (GST; y-axis). In general, a low value yielded less error for both MDD and GST parameters. For each participant, the combination of GST and MDD that produced the lowest gaze prediction error was used.

### Gaze Prediction Error

The model’s AM maximum (see Methods) was used to predict the empirical eye gaze location, which allowed a comparison between model and human data at each timestep throughout the experiment.

A paired-sample t-test revealed that RAGNAROC’s gaze prediction error (*M* = 17.80 dva, *SD* = 2.16 dva) was significantly lower than the baseline formed by randomly selecting any item’s location (*M* = 27.53 dva, *SD* = 2.04 dva), *t*(25) = -45.81, *p* < .001, 95% CI [-10.17, -9.29], as shown in Figure 4. This shows that the model’s predicted attentional dynamics had correspondence to how overt attention was deployed by the participants on a moment-to-moment basis. Furthermore, the average model prediction error was within the near peripheral region^44^. At this level of eccentricity, featural discrimination can be reliably attained with stimulus size adjustments^45,46^.

**Figure 4.**
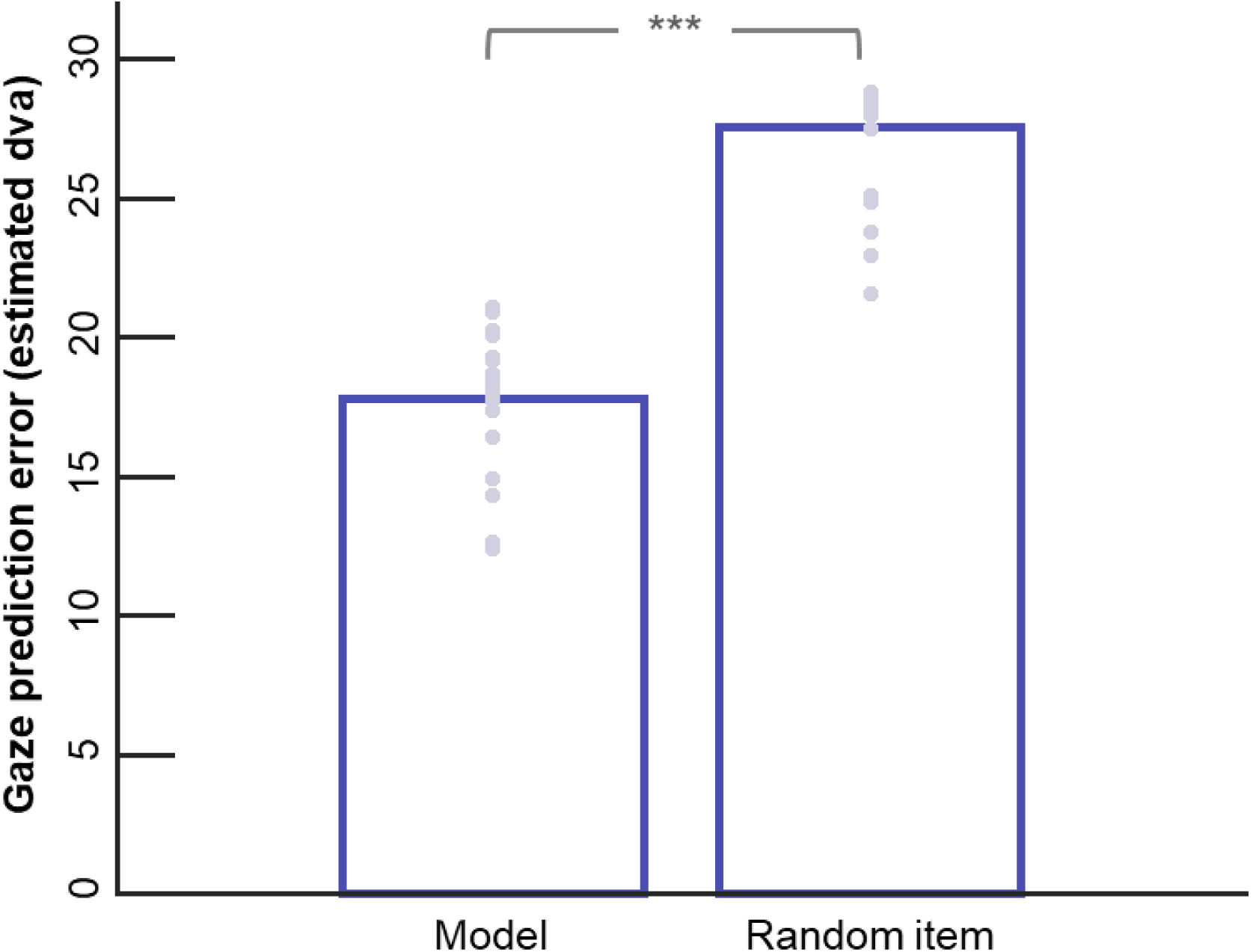
Comparison of mean gaze error as produced by the model versus by random item selection. Bars represent group means. Grey dots represent individual participants.

Although the covert system which RAGNAROC simulates and the overt system which generates eye movements have highly overlapping representations^47,48^, it is evident that they are not identical^49,50^. Using gaze prediction error as metric, such differences between the covert and the overt systems could not be differentiated from limitations in the model’s translatability from the lab to the real world. Moreover, at each timestep, gaze location was a single point, meaning that the simulated priority at only one AM location could be validated. To obtain a more general understanding of the validity of the AM as a whole, additional metrics were considered.

### Item Prediction for Selection Events

As RAGNAROC was parametrized upon desktop paradigms, the model was expected to have an advantage in simulating attention when the available visual field in VR bared a stronger resemblance with a typical desktop display. To this end, time windows immediately before three types of *selection events* were extracted, namely, target fixation, distractor fixation, and the manual response to indicate target selection, allowing the isolation of times when the to-be-selected item was likely already visible. To validate the model, AM priority at the to-be-selected item location was tracked across the pre-selection period and compared with AM priority assigned to a non-selected item. The model was tested on whether it could predict which item was selected next by differentially prioritizing it over other items.

Results are summarized in Figure 5. For target fixations, MUCP (mass univariate test with cluster-based permutation, see Methods) revealed a significantly sized cluster that corresponded to the -390ms to 0ms pre-fixation interval (cluster mass = 239.11, permutation *p* < .0004). In this time interval, the model prediction accuracy (*M* = .57, *SD* = .08) was higher than the baseline formed by random selection (*M* = .47, *SD* = .06). Similarly, a significant cluster was identified prior to distractor fixations (cluster mass = 355.16, permutation *p* < .0004, cluster period: -560ms to 0ms). In this cluster period, the model prediction accuracy (*M* = .12, *SD* = .03) was also higher than the random baseline (*M* = .08, *SD* = .02). As for target shooting, a significantly sized cluster was also found using MUCP, which corresponded to the -1620 ms to - 10 ms interval prior to shot on target (cluster mass = 1271.47, permutation *p* < .0004). The mean model prediction accuracy in this time interval was .62 (*SD* = .08), which was higher than the chance level (*M* = .46, *SD* = .06).

**Figure 5.**
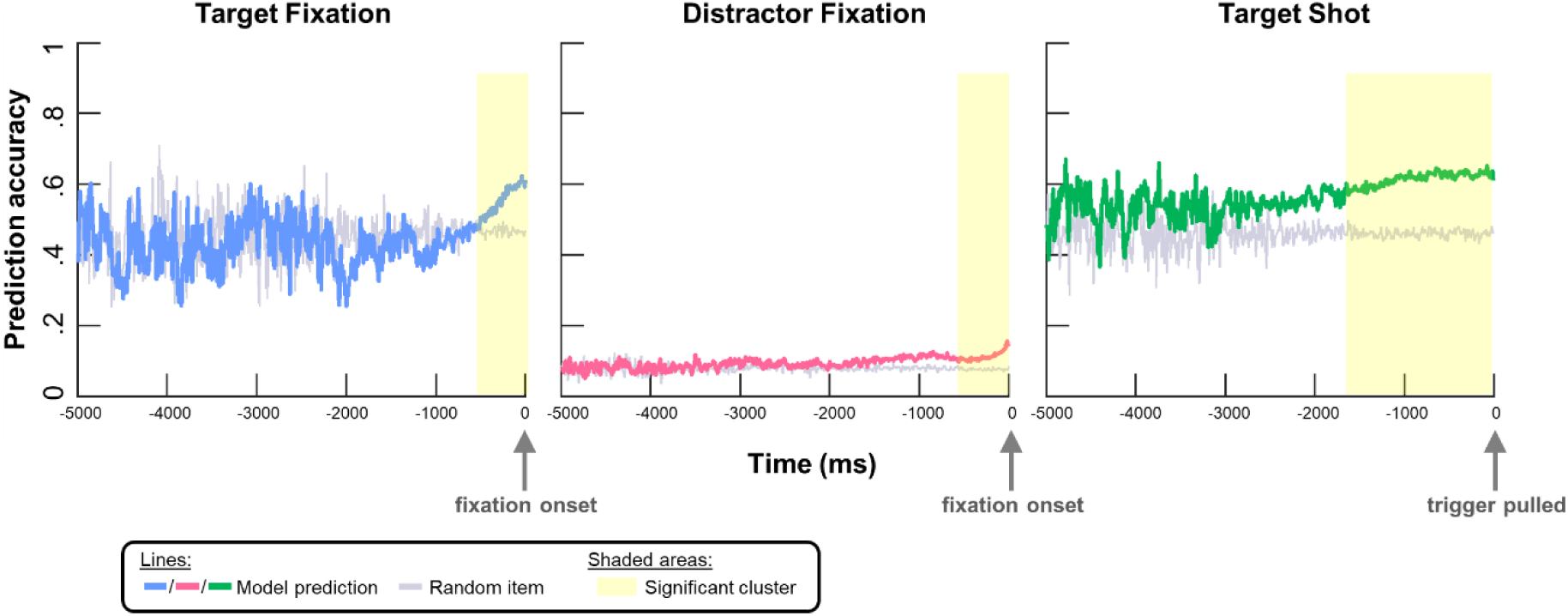
Prediction accuracy of the model (blue / pink / green) and random baseline (grey) over time for target fixation (left graph), distractor fixation (middle graph), and target shot (right graph). Time periods corresponding to significantly sized clusters were highlighted in yellow.

Overall, the model reliably predicted which item was going to be fixated or shot in the immediate future, with the predictivity for shooting emerging earlier and lasting longer compared to that for fixations.

To qualify the model’s simulation success, a set of exploratory analysis was conducted to compare the *properties* of items selected by humans participants versus the model. For each selected item, its *global and local spatial context* was computed (Figure 6). Global spatial context refers to the relative position of the item on the retinotopic map, which was defined as the absolute Euclidean distance between the item and the map center (fovea). The local spatial context was a measure of the crowdedness in the item’s immediate surround (5x5 grid or up to about 11.5 dva from the item, see Methods). Separately for target fixations, distractor fixations, and target shots, all valid pre-fixation / pre-shot timepoints during the corresponding significant cluster periods were considered, where the global and local spatial context scores for all visually available targets (before target fixations and shots) or distractors (before distractor fixations) were computed. For each participant, a median split was performed for both scores, which segregated items into either “central” or “peripheral” with respect to their distance from the retinotopic center, and, independently, either “crowded” or “isolated” in terms their immediate surround. Group statistics of the median thresholds are reported in Table E1.

**Figure 6.**
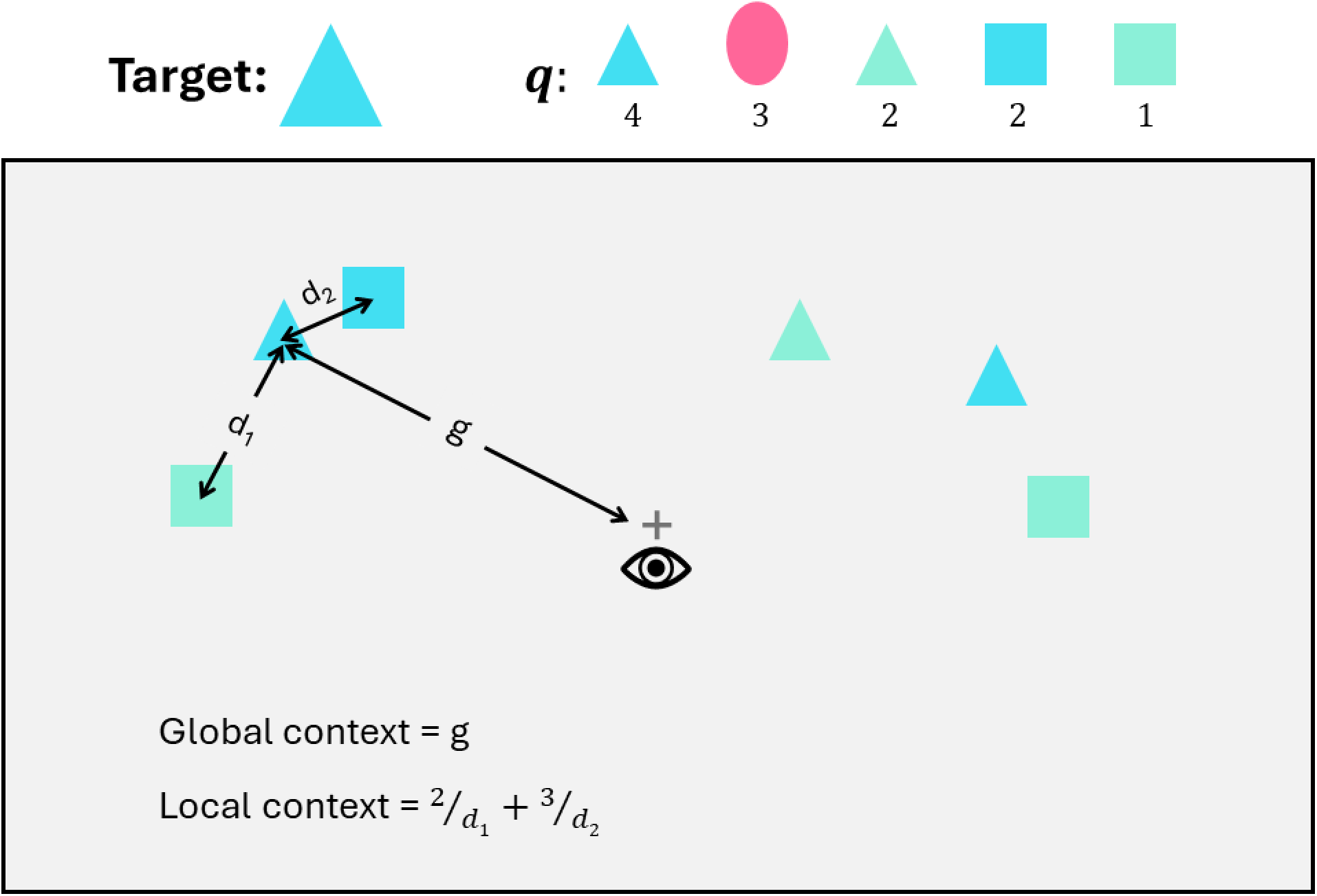
Computation of global and local spatial context. In this example, blue triangles are target, and the spatial context scores for the target on the left visual field are computed. The global context score of the left target is g, which is its distance from the retinotopic center. The local context score is given by the sum of divisions between a surrounding item’s interference score (q) and the distance between the surrounding item and the item-of-interest. The interference score is a value representing the featural overlap with items to be responded to. Therefore, the target shape and the probe have the highest score (q = 4 and q = 3, respectively), distractors with one target-matching feature have the next highest (q = 2) and distractors with no feature overlap with the target have the lowest (q = 1). The surrounding items were defined as items within a 5-by-5 grid centered at the item of interest.

The effects of the item’s global and local spatial context on the probability for it to be fixated or shot next (referred to “selection probability” below) are visualized in Figure 7 (see Table E2 for summary statistics). Regarding empirical selections, a general preference for central items over peripheral items was observed. Moreover, the effect of the item’s local context appeared to depend on its global context, and the pattern of effects varied across selection events (Figure 7A). The significance of the effects of global context, local context, and their interaction on selection probability was tested for each selection event (target fixation, distractor fixation, or target shooting) and selection agent (empirical or RAGNAROC) with a set of Wilcoxon signed-rank tests.

**Figure 7.**
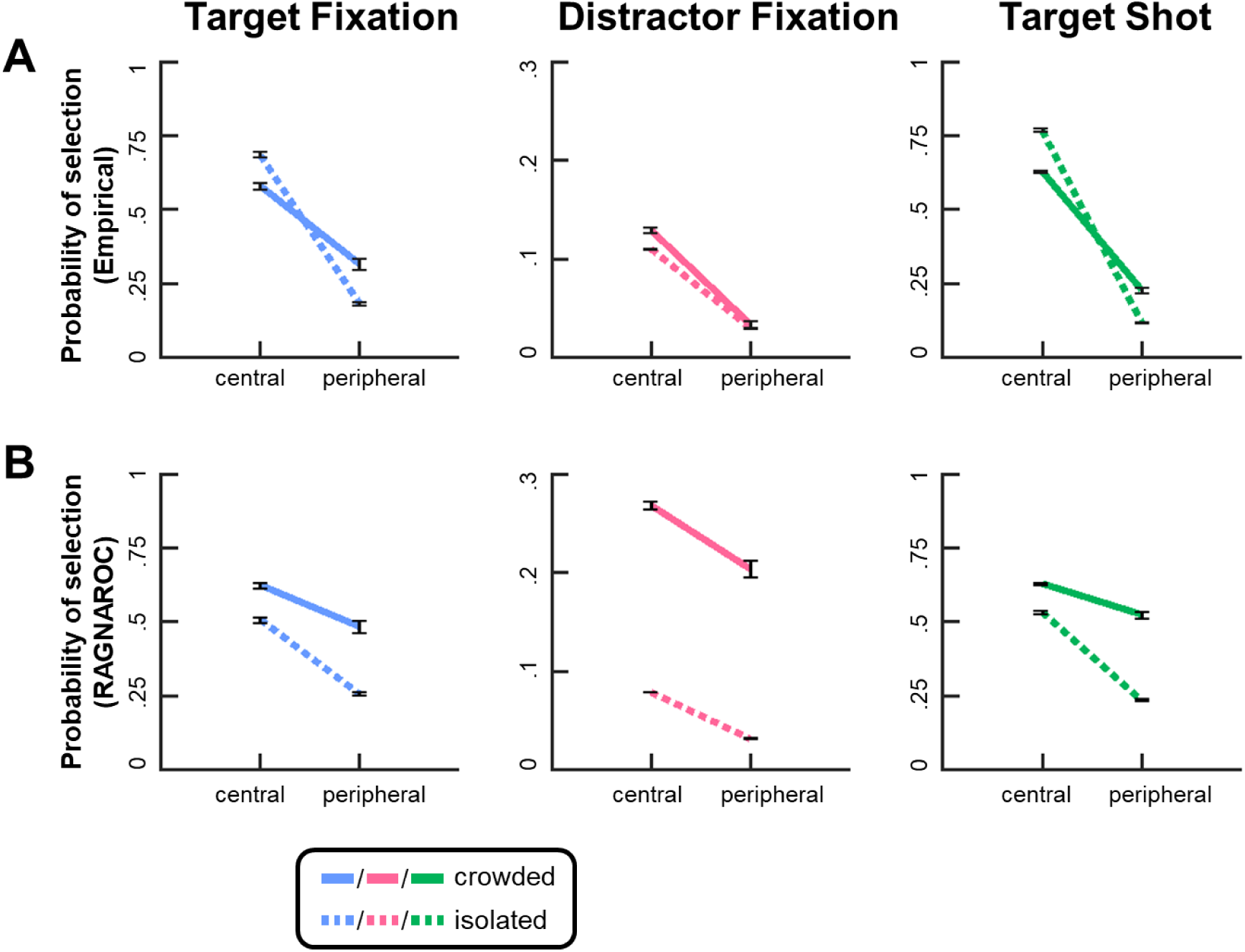
Mean selection probabilities by participants (empirical) and RAGNAROC for items in each spatial context category, separately for each selection event. Error bars represent the means of individual-level standard errors.

#### Selection by participants

In the empirical data (selection by participants), central items were more likely to be selected. For all three types of selection event, the Wilcoxin signed rank was unanimous across participants to show that selection probability was higher for central items compared to peripheral items, *z*s = 4.46, *p*s < .001. Furthermore, target selection was characterized by a crossover interactive pattern where isolated targets were preferred in the central region but crowded targets were preferred in the periphery. The signed rank of each of these effects was unanimous across participants in both cases and similarly for the fixation or shooting of target stimuli (*z*s = ± 4.46, *p*s < .001). For distractors, the signed tests revealed that crowded distractors were preferred in the central region (*z* = 3.29, *p* = .002) but no clear preference between crowded and isolated distractors was found in the periphery (*z* = 1.33, *p* = .365).

#### Selection by model

Consistent with the empirical data, RAGNAROC showed a general preference for items in the central region across selection events, as supported by unanimous signed ranks, *zs* = 4.46, *p*s < .001. However, the crossover interactive pattern between global and local context observed in the empirical selection of targets was not re-produced by RAGNAROC. Instead, crowded items were more likely to be selected by RAGNAROC no matter if they were targets or distractors and occupied a central or peripheral location (target fixation: *z*s = 4.46, *p*s < .001; distractor fixation: *z*s = 4.46, *p*s < .001; target shooting: *z*s > 4.18, *p*s < .001).

### Probe Detection

Probe detection was a secondary task in the experiment that provided another way to validate model predictions beyond a single point. In addition to searching for conjunctively defined targets, participants were instructed to also detect the presence of a sudden onset stimulus that would appear at a pseudorandom timepoint in each trial. Like laboratory paradigms^45^, the reaction time (RT) to this onset stimulus could then be taken as an indicator of the amount of attentional priority assigned at the stimulus’ location. If the model simulated attentional priorities, then it should be able to predict the detection RT.

With a linear mixed effects model where subject-level slope and intercept were included as random effects, a negative association was found between AM priority and probe RT. Specifically, consistent with our hypothesis, higher model priority at the probe location in the first 200 ms of probe presence predicted faster probe RT (*b* = -0.017, *p* < .001, 95% CI [-0.027, - 0.007]). However, data visualization showed that AM priority at the individual-level had an unusual distribution where many participants’ data points were gathered over the same value (Figure E1). This was problematic as it hindered the interpretability of the negative association between AM priority and RT as identified above. The limited range of the priority values observed in some participants was due to the low GST values selected for those participants. A low GST value meant that smaller changes in viewpoint triggered the model to “reset” its upper-level layers (including the AM) which inadvertently limited the time available for activation accumulation and lateral competition, in turn restricting the range of AM priority signals being generated. To verify and quantify this relationship, a correlation test was run between the selected GST and the range of AM priority on the participant level, which showed a strong, significant relationship (*r* = .910, *p* < .001).

To test if relationship between AM priority and probe RT was an artifact of the skewed distribution of model activation across participants, the LME analysis was re-run with model simulations generated higher GST values that were selected to trigger model resets on roughly half of the total timepoints recorded for each participant. This yielded selected GST values that were substantially higher (mean = 16.04 cm, mode = 20.09 cm, SD = 10.08 cm) and thus a larger range in AM priority at the probe location across participants. The same linear mixed effects model specified above was run on these new values and again revealed a negative association between AM priority and probe RT (*b* = -0.0041, *p* = < .001, 95% CI [-0.0066 -0.0017]). In other words, even after configuring the model to allow for a larger variation of AM priority at the probe location, there was still evidence to support that stronger model priority was predictive of faster probe detection (Figure 8).

**Figure 8.**
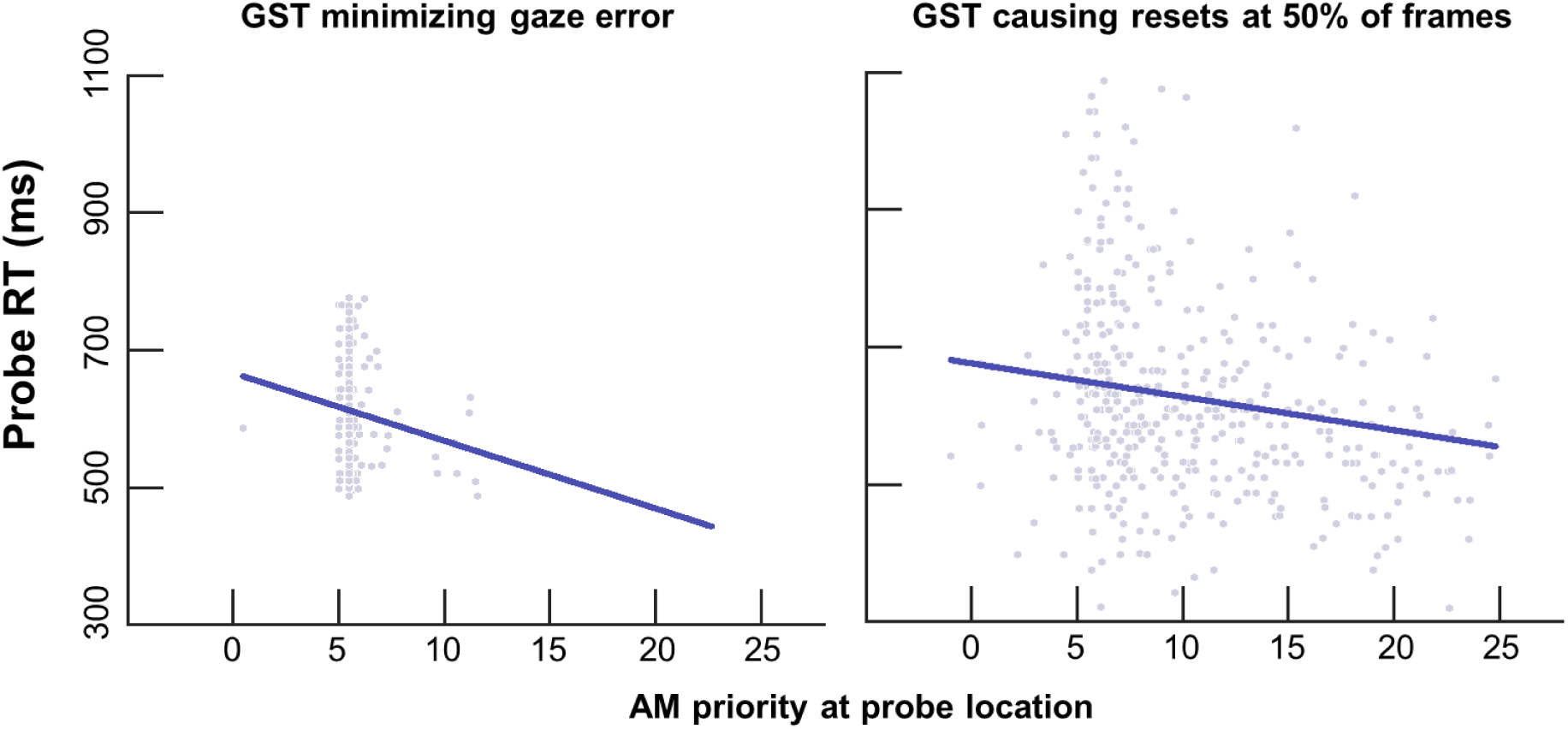
Probe RT plotted against AM priority at the probe’s retinotopic coordinates for the first 200 ms post probe onset. Grey dots represent individual participants. The graph on the left side plots data from model simulations when the GST is set to reduce gaze error per participant (as is used in the rest of the results presented here). The graph on the right side shows data from model simulations when GST is set such that a “reset” of attention map activation only occurs on approximately half of all time steps.

## Discussion

Imposing strict control over stimuli presentation and participant movement has been the prevalent approach underlying the advancement of the field of visual attention. However, while experimental control is necessary, the goal of such work is to inform our understanding of how the human attentional system operates in real-world vision where the environment, and movements of the viewer are far more varied than in the laboratory. In the current work, a foraging task was conducted in VR to move a well-studied conjunctive visual search paradigm into an environment where participants could move their eyes, head, and torso with relatively few constraints. Search behaviors were then compared with predictions generated by RAGNAROC, a model of covert attention that has previously been parametrized on a wide range of behavioral and neurophysiological findings from desktop paradigms and thereby encapsulated a number of key attentional mechanisms that have been identified in the field. Evaluating the model’s predictive capability made it possible to test these mechanisms, collectively, in a more ecologically valid context. In brief, the current work forms a bridge between a wealth of experimental findings in the field and human vision as it arises in the real world.

Together, the results indicated that the neural mechanisms for visual attention simulated in RAGNAROC accounted for a significant portion of variability in human search behavior in an immersive, less constrained environment. Across all the validation metrics, RAGNAROC performed better than chance. Using the priority values of its Attention Map (AM), RAGNAROC was able to predict a participant’s moment-to-moment eye gaze location within near-peripheral regions^44^. In time periods immediately prior to critical selection events, which included item fixation and target shooting, RAGNAROC reliably prioritized the to-be-fixated and to-be-shot item over other same-category items. Finally, RAGNAROC predicted the reaction time performance for a sudden onset probe that participant had to detect as a secondary task. Overall, the results showed promise that the current understanding of visual attention based on desktop paradigms is indeed appropriate to explain behaviors in real-world contexts and demonstrated a well-founded approach to show how such ecological validity can be quantified. Additionally, our results provided new insights into search behaviors exhibited when the search array extended beyond the visual field that may not be predicted by desktop search paradigms.

The results showed that a low gaze shift threshold *(M* = 0.71 cm) produced the lowest gaze error, with most participants fitted with GST of zero, meaning that the AM maximum tracked the moment-to-moment gaze location best when the model’s attentional dynamics were not influenced by retinotopic traces (i.e., activation history on retinotopic maps from previous time steps). This is unexpected as the literature shows strong evidence that attention, by default, operates on retinotopic coordinates and thus retinotopic traces influence attentional decisions^51,52^. Part of this discrepancy was possibly due to the fact that GST was selected by an overt metric (gaze prediction error), which deviated from how the effects of retinotopic trace were typically measured in previous studies (changes in accuracy or RT in responding to a covertly detected stimulus)^51,52^. At the same time, the parameter fitting results might have also reflected that spatiotopic coding was encouraged in the current VR task, which is consistent with evidence showing that retinotopic coding is not exclusive but instead is used conjunctively with spatiotopic or intermediate coding schemes^53,54^, and the use of reference coordinates can vary by task demands^51^. Particularly, spatiotopic coding had been found to be more prominent in dynamic saccade contexts where, like in the current task, frequent eye movements were made^55^. Furthermore, moment-to-moment viewpoint changes were also more drastic than those typically allowed in a laboratory setting, such that there might be a tendency to reduce the use of retinotopic coding to avoid big spatial discrepancies between the attended location and the corresponding item’s updated location. However, our results should not be taken to suggest that large viewpoint changes would lead to a “reset” of the attentional landscape for visual area neurons like RAGNAROC. Instead, the “reset” preformed here was a stopgap measure because RAGNAROC does not have the relevant components to address other possible mechanisms, e.g., predictive remapping^56–58^, involved in managing attentional history across viewpoint changes.

RAGNAROC predicted the to-be-selected item in the time immediately preceding three types of selection events: target fixation, distractor fixation, and target shooting. The empirical selection probability was found to be influenced by the item’s global context (“central” or “peripheral”) and local context (“crowded” or “isolated”), and this selection pattern was partially captured by the model. In human selection, observing a bias for central items was close to mandatory because fixating at an item by definition meant that the item was at the retinotopic map center. Note also that this central bias occurs on retinotopic coordinates, which is not directly comparable to the traditional use of this term, which refers to the tendency to fixate at the central region relative to the object^59^, scene^60^, or even the virtual environment^61^ being viewed. Furthermore, the empirical preferences for crowded versus isolated target items were dependent upon the item’s retinotopic location, as selection was biased towards isolated items near the retinotopic center but crowded items in the periphery. One possible explanation is that there was a switch in decision making criteria when selecting information in the central versus peripheral field of view. In the central region, selection might have been geared to optimize target feature identification, which is generally easier for isolated items (crowding effect^62,63^). Whereas in the periphery, crowded items were instead preferred because the chance of finding a target was simply higher in a crowd (i.e. areas of higher entropy^64^). Focusing target identification on central items could be because identifying a conjunctively defined target requires feature-location binding^43,65^ which is challenging to do in the periphery^66^. Peripheral recognition was also unnecessary in the current task because participants were free to shift their gaze. This differentiation between peripheral and central processing is broadly consistent with the idea that peripheral vision specializes in “looking” whereas central vision specializes in “seeing”^67^, and also not dissimilar from exploitation versus exploration research that shows that search patterns change depending on the context^68^. On the neural-mechanistic level, this could be related to a variation in neural competition mechanics across eccentricities. Specifically, center-surround inhibitory profiles among ventral visual neurons are narrower at the fovea and wider in the periphery^69^, meaning that within the foveal region, crowded representations receive disproportionately strong inhibitory currents which would in turn favor the selection of isolated items.

Differing from the empirical data, RAGNAROC was more likely to select crowded targets regardless of central or peripheral positioning. This was likely attributable to the spatial spread of feedforward excitation across the model’s hierarchy. Specifically, feedforward excitation at each layer in the model is projected with a Gaussian spatial spread such that the priority value of an AM node (top layer) is affected by multiple input neurons on the lower levels of the hierarchy. As a result, an AM node representing a crowded region (i.e., multiple active input nodes) was more likely to gain a higher priority value. As current model selection was estimated by the point of max activation across the attention map, the model was unable to capture differences in selection strategy or mechanisms between central and peripheral areas. To improve simulation accuracy, an alternative selection criterion that maximizes local signal-to-noise ratio could be applied for foveal neurons such that isolated targets would be prioritized. In addition, modeling the variation of inhibitory profile across eccentricities could result in selective suppression of crowded central neural representations and therefore further improve simulation accuracy.

In addition to serving as a vehicle by which to validate a collection of attentional mechanisms, isolated in desktop paradigms, in an immersive, less constrained context, RAGNAROC also has important possible applications in human-centered technologies, which focus on design and systems that complement the natural tendencies of human users^16^. Computer-aided detection (CAD) systems, where salient visual markers are overlaid onto the operator’s visual field to facilitate target detection, are increasingly explored in the military^17–19^ and medical^20,21^ fields. However, these systems can be counterproductive if not calibrated to complement the human attentional system^22^. The quantitative nature of of the model’s attentional deployment predictions turn what information the human has attended to and what information was possibly missed into a machine-readable format, allowing artificial systems to function adaptively to the human user^23^.

Together, the current work showed that the current understanding of many theorized attention mechanisms studied through desktop experimental paradigms is generally applicable to search behaviors in a more naturalistic context. The simulation analysis provided further insights into several important issues that need addressing when transitioning laboratory findings into the real world, including the role of larger-scale self-initiated movements on the reference coordinates of attention and the role of central versus peripheral vision. This is an important step in moving attentional modeling into more ecologically valid environments but also using behavior exhibited in such environments to inform further model development and elucidate cognitive processes that may otherwise go unobserved.

## Methods

### Computational Model: RAGNAROC

RAGNAROC simulates the spatiotemporal deployment of covert attention through a hierarchical structure, where visual information is represented with multiple layers of retinotopically organized artificial neurons. Visual input first travels to the Early Vision (EV) layer, where input locations are encoded. Signals from EV are then modulated by a set of weights that represent bottom-up visual saliency, allowing stimuli with above-threshold saliency to progress in the hierarchy and be represented on the Late Vision (LV) layer. LV output will then be further modulated by a set of weights that represent top-down feature-specific influences such that representational strengths at the top layer, the Attention Map (AM), encompass both bottom-up and top-down information. Simulated attention dynamics emerges from lateral competition among AM neurons and was the basis of RAGNAROC’s predictions in the current study.

The specific mechanism of the AM competition is as follows. The AM is paired with the Inhibitory Gating (IG) map of the same size. As an AM neuron crosses a threshold, IG neurons in the neighboring positions will be excited through a difference-of-Gaussian profile. IG neurons are also excited by LV neurons, and an IG neuron crosses threshold only if it receives excitation from both AM and LV (“AND” gate). An above-threshold IG neuron inhibits the AM neuron at the corresponding location (one-to-one). Together, this means that an excited AM neuron will inhibit its surround but only at locations receiving visual input, simulating empirical results^70^. An AM neuron “wins” the competition when it crosses a high threshold which additionally invokes a one-to-one inhibition to the IG map. This allows the AM neuron to be protected from the surround inhibition centered at other AM neurons, if any. At the same time, the winning AM neuron sends excitatory feedback to EV to achieve recurrent amplification.

RAGNAROC has been parametrized to simulate covert search behaviors and ERP findings from a variety of desktop paradigms. Readers can refer to the original paper^24^ for the full list of simulated phenomena and other computational details of the model.

### VR Task

#### Participants

This study was approved by the U.S. Army Research Laboratory (ARL) Human Research Protection Program (HRPP) and conducted in accordance with the accredited Institutional Review Board (ARL-23-045). All participants signed an IRB approved informed consent form prior to participating in this study. Thirty participants were recruited from Towson University Northeast Campus (11 male, mean age = 25.47). To participate, they had to be at least 18 years old and sufficiently fluent in English to understand the task. After dropouts and the loss of two participants’ data due to technical issues, data from 26 participants were included in the final analysis.

#### Apparatus and Stimuli

The HTC Vive Pro Eye headset was used to record head, eye and weapon movement and to display the virtual environment. The display resolution was 1400 x 1600 per eye with a refresh rate of 90 Hz. The headset was tethered to the experiment computer. The embedded eye tracker had a sampling frequency of 120 Hz with an accuracy of 0.5-1.1 degrees and a trackable field of view of 110 degrees. The eye tracker was calibrated at the start of the experiment with the standard Vive procedure, which included having participants follow a white dot with their eyes from the center of the screen to 6 peripheral locations.

Participants used a polymer Combat Machine CM16 Carbine AEG Airsoft Rifle (approximately 5.25 inches long and 8.5 pounds) to shoot targets. A Vive tracker was mounted on the Picatinny rail of the rifle and connected to the internal trigger mechanism as well as two other buttons that were placed above the trigger on both sides of the rifle (to accommodate right- and left-handed participants). This tracker enabled rifle movements to be integrated into the virtual environment as well as allowing the recording of trigger pulls and button presses. This integrated rifle system was intended to address research questions on the effect of weapon aiming on visual attention. Paradigm details and data analyses related to the weapon aim point will be reported in a separate article.

The virtual environment was developed in the Unreal Engine (Version 4.27). Participants stood in the center of a virtual dome with a radius of 5 meters. The walls of the dome were patterned with a 1/f spatial frequency noise to have a similar spatial power spectra as found in natural images^71^. The search arrays consisted of green pyramids, blue pyramids, blue cubes, and green cubes. The entire array spanned 200° X 130° in front of the participant (Figure 2A). The RGB values were (0, 222, 255) and (255, 97, 51) for blue and green stimuli respectively. All of the search stimuli were 0.18 meters across. In each trial, the search array consisted of 50 stimuli in total including 2 to 8 targets. The number of targets and their locations were randomized per trial. Once during each trial, a probe stimulus was presented. The probe was a red disk (RGB values: 255, 0, 0) with a diameter of 0.18 m and a transparency of .3.

#### Procedure

After fitting the headset and calibrating the eye tracker, participants began the experiment in the center of the virtual dome and were told to find and shoot all of the targets in an array of stimuli as quickly and accurately as possible. Each participant was given one of the 4 stimulus types (green pyramids, blue pyramids, blue cubes, and green cubes) as the target with the other three serving as distractors. Target type was held constant throughout the experiment and was counterbalanced across participants. Participants were also told that during the trial a red dot (the probe) would appear. They were instructed not to shoot the probe but instead to press the button above the trigger as fast as possible when they saw the probe. Participants were told to pull the trigger with their middle finger so that the index finger could rest on the button to facilitate this response.

Each trial began with the participant shooting a green ‘x’ placed directly in front of them. Once shot, this green ‘x’ was replaced by the search array. Unbeknownst to the participant, on every trial one of the targets was randomly selected as the critical target (excluding the targets that were shot first or last). The critical target was not discernably different to the participant but the shooting of it started a time window in which the probe could be presented. Once the critical target was shot, the probe was presented at the next frame in which the following two criteria were met. First, a valid eye position was returned (i.e., the eyes were not closed). Second, the gaze point (the extrapolated position where the gaze vector first intersected with the virtual environment) and the aim point (the extrapolated position where the vector extending from the muzzle of the rifle intersected with the environment) were spaced by 5-8 degrees of visual angle from one another. Once these two criteria were met, the probe would appear at either the gaze point location (where the participant was looking), the aim point location (where the weapon was aimed), or at a neutral location, diametrically opposed to the aim point along an imaginary circle, centered at the gaze point with radius determined by the distance between the gaze and aim point. In this way, neutral probes were presented, on average, at similar eccentricity to the fovea as aim-point probes. Each probe location (gaze / aim / neutral) was equally represented and randomly intermixed within the experiment. Participants were instructed to hit the response button on the side of the rifle as quickly as possible in response to the probe.

Shooting the last target triggered the end of the trial. Participants were presented with a scoreboard that gave them a score on how fast they shot the targets, their shooting accuracy (shots on target divided by the total of number of shots taken in the trial), and how fast they responded to the probe. At the bottom of the scoreboard was a green ‘x’ that they could shoot to begin the next trial. Participants completed 5 practice trials to ensure they could differentiate targets from distractors and to practice aiming and firing the rifle. After the practice trials there were a total of 60 experimental trials, with breaks offered after trial 20 and 40. In total, the experimental session lasted roughly an hour.

#### Saccade Classification

Prior to saccade and fixation detection, eye-tracking data was pre-processed to exclude invalid samples related to eye blinks or other tracker issues during the recording. Specifically, timepoints at which the recorded pupil diameter was below zero or the eye location was out of range (zero or NaN) were marked as invalid. The left eye sample was considered by default but the right eye sample would be used if the left sample was invalid. If both eyes returned invalid samples, the timestep would be marked invalid. For invalid time windows under 50ms, gaze points were interpolated linearly. For invalid time window of 50ms or longer, the time window, along with one timepoint immediately before and after, were excluded from further analyses. The pre-processing steps ensured that the adaptive velocity threshold for saccade detection was not influenced by abrupt velocity changes caused by eye blinks or dropped samples.

Saccade and fixation detection was computed using an adapted version of the EYE-EEG toolbox^72^, which implements the detection algorithm proposed by Engbert and colleagues^73,74^. Eye velocity was first computed separately along the X, Y, and Z axes and mean smoothed over a sliding 5 consecutive sampled time window. For each participant, the median-based standard deviation of velocity along each axis was calculated and a 3D elliptical threshold was set at 6 standard deviations from the mean on each axis. Saccades were then defined as time points in which this 3D velocity threshold was crossed. Saccades classified within 50ms of one another were clustered, with only the largest saccade being kept in each cluster. Fixations were defined as the intervals between saccades.

### Computing Model Predictions

#### Dimensional Reduction

To feed visual information to RAGNAROC, a two-dimensional projection of the moment-to-moment visual input available to the participant was generated. At a sampling rate of 100Hz, participant’s current head orientation was cross referenced with stimulus locations in the VR environment to recreate visual images from the head-centered field of view (FOV). These FOV frames had arbitrary markings that indicated the locations of targets, distractors, the probe, as well as the projections of the participant’s current gaze point. The dimension of a FOV frame was 2560x1600px which represented about 110 degrees of visual angles (dva) widthwise. See Figure 2B for an example.

For each FOV frame, the centroid coordinates were extracted for all stimulus elements in the frame. Before being fed into the model, these stimulus coordinates were resized to fit a 27x17 grid and re-referenced to be centered at the gaze point (retinotopic reference). Each unit distance on the model map therefore corresponded to approximately 4.07 dva. Targets, distractors, and the probe were assigned specific sets of top-down and bottom-up weights with respect to the experimental design (see Table E4 for values).

#### Validation Metrics

The overarching goal of these validation metrics is to assess the consistency between RAGNAROC’s simulated attentional dynamics and the empirical (human) attentional dynamics.

##### Gaze prediction error

To quantify how well AM priority could predict human eye gaze location at each timestep, the moment-to-moment spatial distance between the maximum point on AM and the empirical gaze point was computed. Specifically, the maximum point on AM was defined as the location with the highest simulated attentional priority in a 21-ms sliding window that started at a given timepoint *t*, and gaze prediction error was defined as the absolute Euclidian distance between this maximum point and the empirical gaze point. Note that the empirical gaze point was always the center point as visual information was represented retinotopically. With a within-subjects t-test, we contrasted the model’s gaze prediction error with a random baseline. To compute the random baseline, at each timestep a random item (target, distractor, or probe) was selected. The locations of these random items represented the “prediction” of a null model that was informed of item locations but had no mechanisms to prioritize items of specific types or saliency levels. The average distance between the randomly selected item and the gaze point at the corresponding timestep formed the random baseline. Contrasting the model’s performance with this baseline therefore allowed us to specifically test whether the attentional mechanisms described in the model provided additional advantage for gaze prediction while controlling for the fact that the model was informed of item coordinates. The model was expected to produce lower error than the baseline.

##### Item prediction for upcoming selection event

Time windows leading up to critical selection events, which included target fixation, distractor fixation, and target selection (via shooting), were isolated to analyze how well AM priority could predict which item would be fixated or selected next. In this and the following analysis, AM priority corresponding to input timestep *t* was defined as the two-dimensional map of priority values averaged across the 21-ms time window starting at *t*.

Item fixations were defined as fixations during which at any time, the extrapolated gaze point intersected with a target or distractor stimulus in the virtual world. Moment-to-moment prediction accuracy was computed in the pre-fixation period, separately for target and distractor fixations. The pre-fixation interval was accounted for up to the previous target/distractor fixation offset or 5000 ms, whichever reached first. In addition, a timepoint was considered valid only if the to-be-fixated element as well as at least one other same-category element were in view (e.g., for target fixation analysis, the to-be-fixated target and at least one other target must be in view). The model’s prediction was considered accurate if the to-be-fixated element was assigned the highest AM priority over other same-category elements in view. We tested whether the model outperformed a chance baseline, which was computed by randomly selecting an element from the corresponding category at each valid timepoint.

As for pre-shot analysis, because shooting was only effective on target stimuli (target stimuli disappeared once shot whereas distractor stimuli did not), only target shots and target stimuli were considered. Prediction accuracy was computed in pre-shot intervals for up to the previous trigger pull timepoint or 5000 ms (whichever reached first), and timepoints were included only if the FOV consisted of the to-be-shot target as well as at least one other target. The model’s prediction was considered accurate if it assigned the highest priority to the to-be-shot target among targets in view. The prediction accuracy was expected to be higher than a chance level baseline where a random target was selected at each valid timepoint.

Both fixation and shot prediction accuracy formed event-locked time series data which were then analyzed with a mass univariate analysis followed by a cluster-based permutation test ^75,76^, abbreviated as MUCP analyses. The general method of this analysis are as follows. First, the model accuracy time series and a random baseline were computed for each participant. The mass univariate approach was then applied wherein at each timepoint, the model prediction data was compared with the random baseline with a paired-sample t-test. A timepoint was retained for cluster extraction only if the t-test was statistically significant at alpha = .05, and it had at least two neighboring significant timepoints. Adjacent timepoints were then be grouped to form clusters, with the cluster mass defined as the sum of absolute t-values across the clustered timepoints. The largest cluster mass obtained from the empirical data would then be used in the permutation test.

This was followed by a permutation test with 2,500 iterations. Within an iteration, each individual subject’s data at each timepoint was randomly shuffled between the model and the random conditions. The same mass univariate approach was then applied to the shuffled time series to identify the largest cluster. The simulated largest cluster sizes that were obtained across iterations then formed a null distribution which represented the likely outcome if there was no difference between the model and the random conditions. The permutation p-value was computed as the probability of generating values in the null distribution exceeded the empirical largest cluster size. The test was considered significant if the permutation p-value was less than .05. If the probability was zero, permutation p-value was reported as p < .0004, which corresponds to 1 in 2,500. We reported the largest cluster mass, permutation p, and the time period at which the largest cluster was found, although it should be noted that the permutation test assessed only the significance of the cluster size but not its extent in time ^75,76^.

The global and local spatial context of items selected by human participants and the model were evaluated and compared. Global spatial context was defined as the absolute Euclidean distance between the item and the map center (fovea). Local spatial context was given by the sum of interference scores, 𝑞_𝑖_/𝑑_𝑖_, for the *n* items in the immediate surround of the item-of-interest, 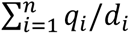. The interference score of each surrounding item 𝑖 was a weighting of the item’s featural relevance and its distance from the central item (*q* = 4 for targets, *q* = 3 for probes, *q* = 2 for distractors with one overlapping feature with targets, and *q* = 1 for distractors with no overlapping features; 𝑑 = absolute Euclidean distance between the surround item and the central item within the local 5x5 grid).

Initially, to evaluate the effects of global context, local context, and their interaction term on selection probability, an ANOVA model was fitted by selection agent (empirical or model) and event (target fixation, distractor fixation, or target shooting). However, the assumption for residual normality was violated in four out of six conditions (Table E3). Therefore, non-parametric tests were instead conducted. Specifically, a set of Wilcoxon signed-rank tests were performed to test the effect of global context on item selection (i.e., comparing selection probability between central and peripheral items), and the interactive pattern between global and local context (i.e., comparing selection probability between crowded and isolated items separately in the central and the peripheral region). When assessing the effect of local context in the crowded and isolated conditions, Bonferroni correction was applied to correct for multiple comparisons. It is also noted here that the typical non-parametric equivalence to the repeated-measures ANOVA is the Friedman test. However, the Friedman test does not test for interaction terms, whereas it was of interest in the current analysis to assess the statistical significance of the interactive pattern observed in the data. This is why we opted to use a set of signed tests to attain statistical analysis for the specific effects of interest.

##### Probe detection

At a pseudorandom timepoint in each trial, an abrupt onset target (probe) appeared at the gaze location, aim location, or a distance-controlled neutral location. This was designed to prompt and compare the underlying attentional priority at those locations, as it was expected that the abrupt-onset probe would be detected faster if it was at a more attended location. The comparison of probe RT among the gaze, aim, and neutral conditions was reported elsewhere^77^. Here, we looked at whether AM priority at the probe location, averaged across the first 200 ms of probe presence, was associated with probe RT. As stronger AM priority in this initial period was expected to indicate more attentional allocation, we predicted a negative relationship where stronger AM priority would be associated with faster probe RT. To test this prediction, we used a linear mixed effects model where RT was predicted with the average model activation as the fixed effect, and subject-level intercepts and slopes as random effects. As probes at gaze and aim point might be modulated by attentional dynamics specific to those locations beyond RAGNAROC’s explanatory power, only the neutral probe condition was included in this analysis. Among the 520 neutral trials, 2 trials were labelled invalid as the participant did not produce a response to the probe. Among the valid trials, trials with RT above or below one standard deviation of the overall group mean (6.95% of valid trials) were further excluded.

#### Individualized Model Parameters

##### Model data delay (MDD)

Visual processing in the brain mandates a time delay between the receiving of visual input and the generation of behavioral outputs such as eye movements and manual selections. In the original version of the model, RAGNAROC had a static time delay of 100 ms between the onset time of visual information and the time at which EV neurons received excitation. This delay was set to simulate the delay between the timepoint when visual information reaches the retina and the timepoint when signals are transmitted to the primary visual cortex^78^. However, this parameter was set when the model focused on simulating discrete episodes of attentional responses for static visual displays presented on a trial-by-trial basis, where the simulated behavioral outputs, despite delayed, could be unambiguously anchored at one input timestep. In contrast, the model was now used to predict moment-to-moment behaviors where the viewpoint (and thus relative retinotopic locations of stimuli) changed from frame to frame, such that the correspondence between the “output” timestep at which an attentional behavior was recorded (e.g., target fixation onset) and the “input” timestep which marked the visual input driving that behavior was ambiguous. Therefore, instead of inserting a delay within the model runtime, the time allotted to visual information processing in the brain was instead accounted for by inserting a time gap between the generation timestep and the validation timestep of model predictions. Specifically, the model data delay (MDD) characterized the variable time delay between the input timestep *t,* from which model predictions were generated (across a 21-ms sliding window starting at *t*), and the prediction timestep *t* + MDD, which was the timestep where behaviors were read and compared against the model predictions.

##### Gaze shift threshold (GST)

To simulate visual input into the model, discrete FOV images were inputted at each timepoint. However, it is important to note that their origin is continuous, i.e., they represent auto-correlated visual information as viewpoint changes across time. This continuity in the model is represented in the activation of upper-level layers. Namely, input data at each sampled timepoint replaced the EV layer but the upper-level layers’ (LV and AM) dynamics continue developing. This was how the attentional landscape was exploratorily simulated in applying RAGNAROC to video viewing ^24^. However, in the current task, the FOV input critically differs from video data as changes of input across timepoints were caused by self-initiated, instead of external movements. Although much research exists on this topic, the exact mechanisms which facilitate visual stability across viewpoint changes remain unclear ^79–82^. For example, while the visual cortex has the capability to encode information in multiple reference coordinates to facilitate the stability of attended locations across gaze shifts ^56,58,83^, the engagement of alternate reference coordinates aside from retinotopy does not appear to be mandatory ^51,52,84^. To complicate the issue, head movements are essential for navigating in three-dimensional environments but they also affect how gaze shifts are deployed ^85,86^, as well as the representational nature of visual information ^54,87^. However, far less is known about mechanisms to maintain visual stability across head or body movements.

RAGNAROC does not have specific mechanisms to account for visual stability and exclusively uses retinotopic maps. The consequence of allowing upper-level dynamics to continually evolve across timesteps is that it essentially creates lingering “retinotopic traces” that can influence attentional decisions. For example, attentional priority has been shown to be temporarily maintained at the retinotopic position that previously consisted of a search target even if it no longer corresponded to any items after a saccade^51,52^. However, the role of retinotopic traces is less clear in time periods around head or body movements. Head or body movements can lead to drastic changes in viewpoint such that there is little to no overlap in the content of the visual field before versus after the movement. In those cases, maintaining visual stability likely has less to do with integration of retinotopic information but perhaps draws more heavily on visual short-term memory, which has been found to play a more critical role in visual search when the search target is not in view ^88^.

Because of the variation in the extent and source of viewpoint changes in the current experiment, we allowed an element of flexibility for the model to handle continuous input differently depending on the amplitude of viewpoint change. First, the amplitude of viewpoint change was estimated by the absolute Euclidian distance between two consecutive gaze points in world coordinates. Next, an amplitude threshold was set, referred to as the Gaze Shift Threshold (GST). When the amplitude of the viewpoint change was below the GST, only the EV layer was reset (as previously described). However, when the GST was crossed, the upper-level layers of the model and the feedback circuit dynamics were reset in addition to the EV layer. The consequence of this was that retinotopic trace would be eliminated from one timepoint to the next, if the viewpoint change was sufficiently large between timepoints.

##### Parameter search and selection

Parameter search was performed across MDD values ranging from 0 to 960ms with 60ms increments and GST values of 0 (such that the model completely resets at every timepoint), and the exponential *e^x^* for values *x* ranging from 0 to 6 in 1-unit increment. To reduce the chance of overfitting and to enhance computational efficiency, parameter search was completed with a subsample of 10 trials and within each trial, up to 1000 independent model runtimes or half of the total number of model runtimes. The combination of MDD and GST yielding the lowest mean gaze prediction error within this subsample was applied to the full sample of trials and model runtimes to compute the rest of the predictivity metrics.

## Extended Data

**Table E1.**
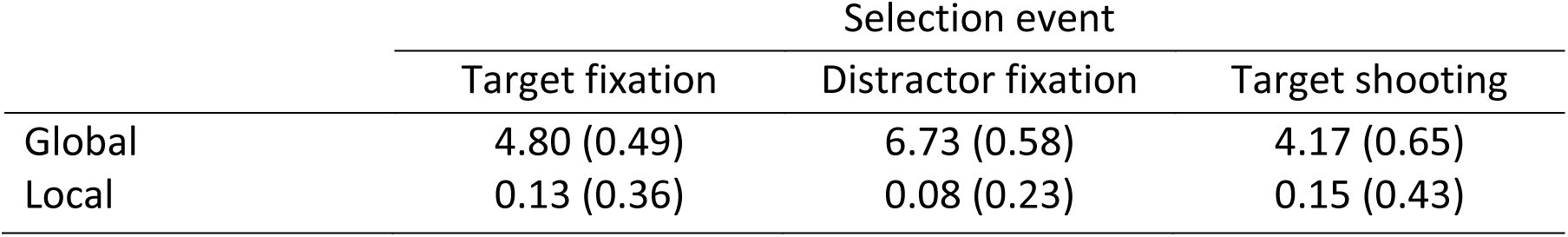
Median threshold values of items considered between each selection event. Median threshold for global context categorized items into either “central” (below median) or “peripheral” (above median). Median threshold for local context categorized items into either “isolated” (below median) or “crowded” (above median).

**Table E2.**
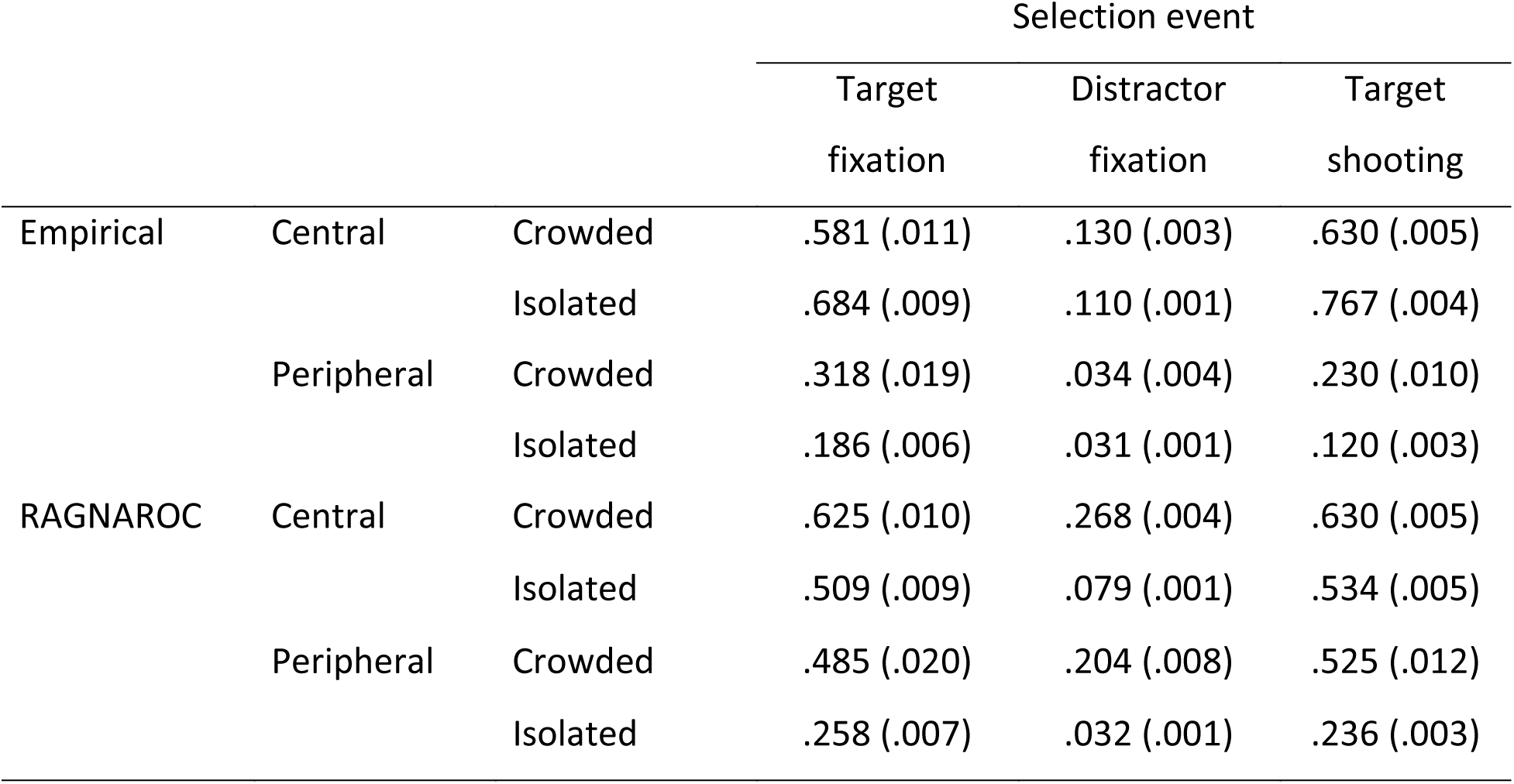
Selection probabilities by participants (empirical) and RAGNAROC for items in each spatial context category, separately for each selection event. Mean probabilities are reported with mean of individual-level standard errors in parentheses.

|  |  |  | Selection event |  |  |
| --- | --- | --- | --- | --- | --- |
|  |  |  | Target fixation | Distractor fixation | Target shooting |
| Empirical | Central | Crowded | .581 (.011) | .130 (.003) | .630 (.005) |
|  |  | Isolated | .684 (.009) | .110 (.001) | .767 (.004) |
|  | Peripheral | Crowded | .318 (.019) | .034 (.004) | .230 (.010) |
|  |  | Isolated | .186 (.006) | .031 (.001) | .120 (.003) |
| RAGNAROC | Central | Crowded | .625 (.010) | .268 (.004) | .630 (.005) |
|  |  | Isolated | .509 (.009) | .079 (.001) | .534 (.005) |
|  | Peripheral | Crowded | .485 (.020) | .204 (.008) | .525 (.012) |
|  |  | Isolated | .258 (.007) | .032 (.001) | .236 (.003) |

**Table E3.** Normality assumption test statistics. For each selection event and separately for empirical and RAGNAROC, a repeated-measures ANOVA model was fitted where global context, local context, and their interaction term were used to predict selection probability. The residual values were then tested for normality using the Shapiro-Wilk test. The assumption of residual normality was considered violated if the Shapiro-Wilk test returned significant, which was the case in four out of six situations.

|  |  | Target fixation | Distractor fixation | Target shooting |
| --- | --- | --- | --- | --- |
| Empirical | W | .986 | .949 | .972 |
|  | p | .296 | < .001 | .026 |
| RAGNAROC | W | .982 | .904 | .934 |
|  | p | .156 | < .001 | < .001 |

**Table E4.** Saliency and relevance weights assigned to each type of stimuli. As the target was conjunctively defined, distractors were differentiated into “match” distractors which had either the color or the shape target feature and “mismatch” distractors which had a different color and a different shape compared to the target.

|  | Saliency weight | Relevance weight |
| --- | --- | --- |
| Target | .2 | .5 |
| Distractors (match) | .2 | .1 |
| Distractors (mismatch) | .2 | .05 |
| Probe | .3 | .5 |

**Figure E1.**
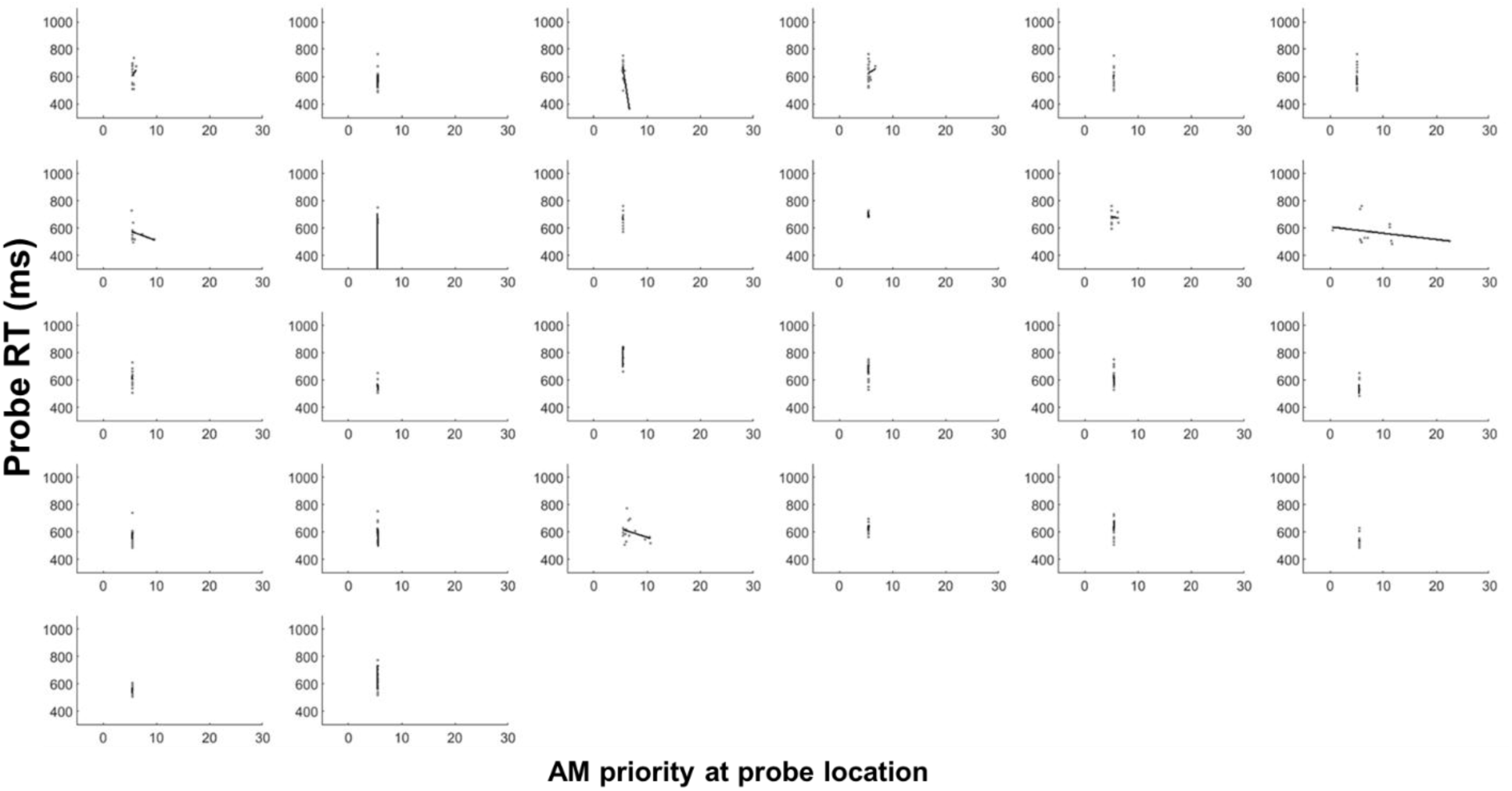
Individual-level relationship between AM priority and probe RT with low GST. AM priority clustered over the same value for many participants because when setting the GST to minimize gaze prediction error, a low GST value was selected for those participants, leading to frequent model resets and limiting the opportunity for AM priority to accumulate. The fitted line for each participant was generated from a linear regression model using AM priority to predict probe RT. Group-level data using this GST setting is visualized in Figure 9 (left).

